# Caffeine consumption during pregnancy accelerates the development of cognitive deficits in offspring in a model of tauopathy

**DOI:** 10.1101/710749

**Authors:** Stefania Zappettini, Emilie Faivre, Antoine Ghestem, Sébastien Carrier, Luc Buée, David Blum, Monique Esclapez, Christophe Bernard

**Affiliations:** Aix Marseille Univ, INSERM, INS, Institut de Neurosciences de Systèmes, Marseille, France; Univ. Lille, Inserm, CHU Lille, UMR-S 1172 - JPArc, LabEx DISTALZ, F-59000 Lille, France

**Keywords:** Caffeine, development, Alzheimer, tauopathy, hippocampus, memory, learning, synaptic currents

## Abstract

Psychoactive drugs used during pregnancy can affect the development of the brain of offspring, directly triggering neurological disorders or increasing the risk for their occurrence. Caffeine is the most widely consumed psychoactive drug, including during pregnancy. In Wild type mice, early life exposure to caffeine renders offspring more susceptible to seizures. Here, we tested the long-term consequences of early life exposure to caffeine in THY-Tau22 transgenic mice, a model of Alzheimer’s disease-like Tau pathology. Caffeine exposed mutant offspring developed cognitive earlier than water treated mutants. Electrophysiological recordings of hippocampal CA1 pyramidal cells *in vitro* revealed that early life exposure to caffeine changed the way the glutamatergic and GABAergic drives were modified by the Tau pathology. We conclude that early-life exposure to caffeine affects the Tau phenotype and we suggest that caffeine exposure during pregnancy may constitute a risk-factor for early onset of Alzheimer’s disease-like pathology.

## Introduction

Adult brain structure is primarily established in early life (Lenroot and Giedd, 2006). Disturbances in anatomical (number and connectivity of neurons) and functional (ability to engage alternative brain networks) development can interfere with crucial processes, including cell proliferation, neuronal migration (Crandall et al., 2004) and post-migration cortical connectivity (Guerrini and Dobyns, 2014). Disturbances can be genetic (e.g. a mutation) and environmental (e.g. maternal separation), resulting in increased risk to develop pathologies during early life such as cortical malformations, developmental delay, epilepsy, and/or autism (Barkovich et al., 2012; Guerrini and Dobyns, 2014), as well as during adulthood/aging, leading to dementia includingsuch as Alzheimer’s Disease (AD)-related pathology (Borenstein et al., 2006; Whalley et al., 2006; Stern, 2012; Seifan et al., 2015).

Neuropathologically, AD is defined by the extracellular accumulation of amyloid beta (Aß) peptides into amyloid plaques, and the presence of intraneuronal fibrillar aggregates of hyper- and abnormally-phosphorylated tau proteins (Masters et al., 1985; Sergeant et al., 2008). Tau pathology is observed early in the brain stem and entorhinal cortex (Braak et al., 2011) and its progression from entorhinal cortex, to the hippocampus, and finally neocortex corresponds to the progression of the symptoms in AD (Duyckaerts et al., 1997; Grober et al., 1999) supporting a pivotal role of Tau pathology in AD-related memory impairments. Various genetic and environmental risk factors are associated with dementia and/or AD (Reitz et al., 2011). Most of these environmental and lifestyle-related factors also impact AD lesions, and in particular Tau pathology. For instance, physical exercise (Belarbi et al., 2011), anesthetics (Le Freche et al., 2012; Whittington et al., 2013), or obesity/diabetes (Leboucher et al., 2013; Papon et al., 2013) modulate Tau pathology and associated memory disturbances.

Among the numerous existing environmental risk factors, those occurring during pregnancy and lactation can have important functional consequences. Exposure to psychoactive substances *in utero* can alter fetal brain development, leading to pathological states later in life for the offspring, including psychiatric disorders (Marroun, 2015; Skorput, 2015). Caffeine is the most frequently consumed psychoactive substance, including during pregnancy (Mandel, 2002; Greenwood et al., 2014). In mice, caffeine exposure during pregnancy and until weaning delays the migration and integration of GABA neurons, enhances seizure susceptibility, as well as alters brain rhythms and hippocampus-dependent memory function in the offspring (Silva et al., 2013; Fazeli et al., 2017). Although it is difficult to generalize rodent studies to humans, a study in mother-child pairs showed an association between caffeine exposure during pregnancy and impaired cognitive development (Galéra et al., 2015). Guidelines for pregnant women recommend to limit the amount of caffeine consumption to 200-300 mg/kg (Gynecologists, 2010). Whether early life exposure to caffeine may prime exposed offsprings to the development of neurodegenerative disorders later in life remains unknown. In the present study, we specifically aimed at determining whether Tau pathology related pathological traits would appear sooner in animals exposed to caffeine during brain development. To address this question, we evaluated the effects of early life caffeine exposure in offspring of the THY-Tau22 transgenic mouse model that progressively develops AD-like hippocampal Tau pathology, with ongoing deficits at 6-8 months of age and a full pathology and memory impairments occurring at 12 months of age (Van der Jeugd et al., 2013).

## MATERIALS AND METHODS

### Animals

Male mice were housed in standard mouse cages under conventional laboratory conditions (12 h/12 h dark-light cycle, constant temperature, constant humidity, and food and water *ad libitum*.). Animal care and experimental procedures were conducted in accordance with institutional guidelines and with the European Communities Council Directive 86/609/EEC and were approved by the Aix-Marseille University Chancellor’s Animal Research Committee, Marseille (France). Since age is a highly important factor in studies related to Tauophaties as a major determinant of phenotype and disease progression, experiments were performed in 8 and 12 month-old mice.

### Caffeine treatment

This study is designed to mimic the situation in pregnant women consuming caffeine before and during their entire pregnancy until full-term birth, both in terms of time span of caffeine exposure and caffeine concentration. In the caffeine groups, female mice received caffeine via their drinking water. Caffeine treatment was started two weeks before mating — reflecting the situation in women who usually do not start consuming caffeine with the onset of pregnancy but did consume caffeine beforehand — and was continued up until postnatal day 15. This end point of the treatment period was chosen based on the assumption that, from a developmental point of view, term birth in humans (= end of gestational week 40) is approximately comparable to postnatal days 12/13 in rodents (Finlay, 2001; Romijn et al., 1991). Therefore, caffeine exposure during the entire pregnancy up until postnatal day 15 in mice would approximately reflect pregnancy until term birth in humans. Caffeine powder was dissolved in tap water at a concentration of 0.3 g/l and given as drinking water to the dam. This concentration was chosen based on previous findings that exposure to 0.3g/l caffeine via drinking water results in a plasma caffeine concentration in rat dams that is similar to that found in the blood of humans drinking three to four cups of coffee per day (Adén et al., 2000) and in a plasma concentration of caffeine in rat pups at P7 similar to that found in the umbilical cord of human neonates whose mothers consume moderate amounts of coffee (up to three cups per day) (Björklund et al., 2008). In a previous study, we confirmed that this caffeine concentration in mice results in a serum concentration comparable to the one previously reported in rats (Silva et al., 2013). Also, we showed that this treatment regimen only affects maternal water intake on the first day of treatment, but not in the following time span (Silva et al., 2013), suggesting that two weeks of treatment prior to mating should be enough to reach stable caffeine concentrations in the dam at the onset of pregnancy.

From postnatal day 15 on, dams and offspring received pure drinking water. The water groups was never exposed to caffeine and received pure drinking water at all times.

### Electrophysiology in hippocampal slices

The barrage of synaptic inputs received by neurons can characterize the functional state of neuronal networks. To quantify the excitatory and inhibitory inputs received by CA1 hippocampal pyramidal cells, we measured the frequency and amplitude of spontaneous and miniature excitatory and inhibitory postsynaptic currents in acute slices from wild type, wild type caffeine, Tau water and Tau caffeine mice.

Number of animals per group: n= 8 cells, 8 slices, from 5 Wild type water mice vs. n= 8 cells, 8 slices, from 5 Wild type caffeine-exposed mice, vs n= 9 cells, 9 slices, from 5 Tau water mice vs. n= 9 cells, 9 slices, from 5 Tau caffeine-exposed mice (8 months) and n= 8 cells, 8 slices, from 5 Wild type water mice vs. n= 8 cells, 8 slices, from 5 Wild type caffeine-exposed mice, vs n= 7 cells, 7 slices, from 4 Tau water mice vs. n= 9 cells, 9 slices, from 5 Tau caffeine-exposed mice (12 months). Transverse cortical slices (350 μm) were prepared with a vibroslicer Leica VT 1200S in a cold (lower than 4 °C) cutting solution containing 140.0 mM potassium gluconate, 10.0 mM HEPES, 15.0 mM sodium gluconate, 0.2 mM EGTA, 4.0 mM NaCl, pH 7.2. After 20 minutes recovery in a preincubation solution (110 mM Choline chloride, 2.5 mM KCl, 1.25 mM NaH_2_PO_4_, 10 mM MgCl_2_, 0.5 mM CaCl_2_, 25 mM NaHCO_3_, 10 mM D-glucose, 5 mM sodium pyruvate equilibrated with 5% CO_2_ in 95% O_2_ at room temperature), slices were perfused for at least one hour with aCSF containing 126.0 mM NaCl, 25.0 mM NaHCO_3_, 10.0 mM D-glucose, 3.5 mM KCl, 2.0 mM CaCl_2_, 1.3 mM MgCl_2_.6H_2_O and 1.2 mM NaH_2_PO_4_ equilibrated with 5% CO_2_ in 95% O_2_ at room temperature and transferred to a chamber containing the same aCSF, kept at a temperature between 33 °C and 35 °C. Cells were recorded under visual control (Nikon FN1 microscope – Scientifica Patch Star manipulators) with an Axopatch700B amplifier and Digidata 1322 interface (Axon Instruments). Currents were recorded using an internal pipette solution of 120.0 mM CsGluconate, 20.0 mM CsCl, 1.1 mM EGTA, 0.1 mM CaCl_2_.2H_2_O, 10.0 mM HEPES, 2.0 mM Mg-ATP, 0.4 mM Na-GTP, 2 mM MgCl_2_.6H_2_O, CsOH.H_2_O to adjust pH (pH 7.3, 280 mOsM). Inhibitory Post-Synaptic Currents (IPSCs) were recorded at a holding potential of +10mV, the reversal potential for glutamatergic events; Excitatory PSCs (EPSCs) were recorded at −60 mV, the reversal potential for GABAergic events (Cossart et al. 2001). We also measured miniature excitatory postsynaptic currents (mEPSCs) and miniature inhibitory postsynaptic currents (mIPSCs) in acute slices from the four groups at 8 and 12 months, after the addition of TTX (1 µM). Average resistance series (RS) in neurons from 8 months wild type, wild type caffeine, Tau water and Tau caffeine mice were 35.3 ± 3.6 MΩ, 34.91 ± 2.6 MΩ, 23.58 ± 2.8 MΩ and 20.70 ± 3.1 MΩ, respectively (*p< 0.05Wild type vs Tau water and Tau caffeine, p<0.01 Wild type caffeine vs Tau caffeine*); in neurons from 12 months Wild type, Wild type caffeine, Tau water and Tau caffeine mice were 31.0 ± 2.9 MΩ, 22.88 ± 3.0 MΩ, 27.32 ± 3.6 MΩ and 24.34 ± 3.0 MΩ, respectively (*p> 0.05*). Although RS is in the range of that found in old control animals, we do not know why it is lower in Tau mice. When the RS changed during the recording by more than 20%, the recording was terminated. Clampfit 10.2 and Minianalysis software (Synaptosoft) were used to analyze synaptic events. All recorded cells were filled with biocytin for *post-hoc* morphological identification, which was performed according to a previously described protocol (Esclapez et al., 1999). The two-sample Kolmogorov-Smirnov test was used for statistical analyses between groups. The level of significance was set at p<0.05.

### Barnes Maze

Barnes maze was used to compare cognitive deficits in learning and memory of 8- and 12-months Tau caffeine (n= 13 and 10), TAU water (n= 11 and 4) mice to the Wild type caffeine (n=13 and 10) and Wild type (n=11 and 8) groups. The maze was made from a circular, 13-mm thick, white PVC slab with a diameter of 1.2 m. Twenty holes with a diameter of 44.4 mm were made on the perimeter at a distance of 25.4 mm from the edge. This circular platform was then mounted on top of a rotating stool, 0.89 m above the ground and balanced. The escape cage was made by using a mouse cage and assembling a platform and ramp 25.4 mm below the surface of the maze. The platform, made of a square petridish, and ramp, made of laminated cardboard, were made out of plastic to be easily cleanable with 70% ethanol. The outside of the walls of the cage was covered with black paper to make the inside of the cage dark and thus attractive to the mice. The maze was placed in the center of a dedicated room and two 120 W lights were placed on the edges of the room facing towards the ceiling about 3/4 of the way up from the floor and about 3–5 feet away from the maze. Eight simple colored-paper shapes (squares, triangles, circles) were mounted around the room as visual cues, in addition to the asymmetry of the room itself. After testing each mouse, the cleaning of the quadrant of the maze around the target hole was alternated with cleaning the whole maze, using 70% ethanol. The maze was rotated clockwise after every 3 mice to avoid intra-maze odor or visual cues. All sessions were recorded using COP Security Monochrome CCD Camera (Model 15-CC20) and MyTV/x software (Eskape Labs).

The animals interacted with the Barnes maze in three phases: habituation (1 day), training (3 days), probes (2 days: one day after one week from the last trial day = Probe 1; one day after two weeks from the last trial day = Probe 2). Before starting each experiment, mice were acclimated to the testing room for 1 h. Then all mice (n = 2–4) from one cage were placed in individual holding cages where they remained until the end of their testing sessions. Holding cages were used during the experiment to control for potential artifacts that could result from housing some mice only two per cage and remained alone while the other mouse was being tested, compared to other mice that were housed four per cage and therefore were never left on their own. Additionally, using holding cages prevented potential influence by mice that had already completed the test on the mice waiting for their turn. After all mice from one home cage completed testing for the day, they were placed back in their home cage together, the holding cages were cleaned, and the next set of mice was separated into individual holding cages. On the habituation day, the mice were placed in the center of the maze underneath a clear 3,500-ml glass beaker for 30 s while white noise was played through a sound system. Then, the mice were guided slowly by moving the glass beaker, over 10–15 s to the target hole that leads to the escape cage. The mice were then given 3 min to independently enter through the target hole into the escape cage. If they did not enter on their own during that time, they were nudged with the beaker to enter. Getting the mice to enter the escape cages is key in “showing” them that the escape cage exists and gives them practice in stepping down to the platform in the cage. The mice were allowed to stay in the escape cage for 1 min before being returned to the holding cage. Once all animals had completed the 1-session habituation, they were all returned to their home cage. In the training phase, mice were placed inside an opaque cardboard cylinder, 25.4 cm tall and 17.8 cm in diameter, in the center of the Barnes maze for 15 s. This allowed the mice to be facing a random direction when the cylinder was lifted and the trial began. At the end of the holding period, a buzzer was turned on, the cylinder was removed, and the mice were allowed to explore the maze for 2 min. If a mouse found the target hole and entered the escape cage during that time, the end-point of the trial, it was allowed to stay in the escape cage for 1 min before being returned to the holding cage. If it did not find the target hole, the mouse was guided to the escape hole using the glass beaker and allowed to enter the escape cage independently. If it did not enter the escape cage within 3 min, it was nudged with the beaker until it did. If a mouse still did not enter the escape cage after 1 min of nudging, it was picked up and manually put on the platform in the escape cage. Then it was allowed 1 min inside the escape cage before being returned to the holding cage. In all cases, the buzzer was turned off once the mouse entered the escape cage. This process typically took 5–7 min per mouse and was done with four mice at a time, providing a 20–30 min inter-trial interval. The total number of trials used was 3 trials on training day 1, 2 and 3. During the training phase, measures of primary latency were recorded. Primary latency was defined as the time to identify the target hole the first time, as mice did not always enter the hole upon first identifying it. Parameters were assessed by blinded observers. About 70% of the measures were randomly reassessed by a second blinded observer to identify potential inaccuracies. Differences between the two observers were insignificant in all cases. In all the cases in which two observers scored the raw data, their scores were averaged. On the probe day, 7 and 14 days after the last training day, the escape cage was removed, mice were placed inside the opaque cylinder in the center of the maze for 15 s, the buzzer was turned on and the cylinder removed. Each mouse was given 2 min to explore the maze, at the end of which, the buzzer was turned off and the mouse was returned to its holding cage. During the probe phase, measures of time spent per quadrant and holes searched (HS) per quadrant were recorded. HS was defined as nose pokes and head deflections over any hole. Primary HS was defined as the HS before identifying the target hole for the first time. For these analyses, the maze was divided into quadrants consisting of 5 holes with the target hole (goal) in the center of the target quadrant (goal zone). The other quadrants going clockwise from the goal zone were labeled: right, opposite, and left zone; all the data collected out of the goal zone are expressed as “other zone”. Data are shown as means 6 standard error of the mean (SEM). Two-way repeated measures ANOVA followed by Bonferroni post-hoc analysis were used for probe day and training trials data, respectively. The level of significance was set at p<0.01.

### Biochemical and molecular evaluations

Animals were sacrificed at 6 months of age by cervical dislocation, brains harvested, left and right hippocampi dissected out using a coronal acrylic slicer (Delta Microscopies) at 4°C and stored at −80°C for biochemical and mRNA analyses. For Tau biochemistry, tissue was homogenized in 200μl Tris buffer (pH 7.4) containing 10% sucrose and protease inhibitors (Complete; Roche Diagnostics), sonicated and kept at −80°C until use. Protein amounts were quantified using the BCA assay (Pierce), and samples diluted with lithium dodecyl sulphate buffer (2) supplemented with reducing agents (Invitrogen) and then separated on 4–12% NuPAGE Novex gels (Invitrogen). Proteins were transferred to nitrocellulose membranes, which were then saturated with 5% non-fat dried milk or 5% bovine serum albumin in TNT (Tris 15mM pH 8, NaCl 140mM, 0.05% Tween) and incubated at 4°C for 24 hours with the primary antibodies. All primary antibodies used are described in Table 1. Appropriate HRP-conjugated secondary antibodies (anti-mouse PI-2000 et anti-rabbit PI-1000, Vector Laboratories) were incubated for 1 hour at room temperature and signals were visualized using chemoluminescence kits (ECL, Amersham Bioscience) and a LAS4000 imaging system (Fujifilm). Results were normalized to actin or GAPDH and quantifications were performed using ImageJ software (Scion Software).

**Table 1.**

| Name | Abbreviation | Epitope | Tissue | Type | Origin | Provider | WB |
| --- | --- | --- | --- | --- | --- | --- | --- |
| Anti-Phospho-Tau (AT270) | pT181 | pT181 | M | mono | mouse | Invitrogen #MN1050 | 1/1 000 |
| Anti-Tau 1, clone PC1C6 | Tau1 | Non-phospho-S195, 198, 199, 202 | M | mono | mouse | Millipore #MAB3420 | 1/10 000 |
| Anti-Phospho-Tau (AT100) | pT212/S214 | pT212/S214 | M | mono | mouse | Invitrogen #MN1060 | 1/1 000 |
| Anti-Phospho-Tau (Ser262) | pS262 | pS262 | M | poly | rabbit | Invitrogen #44-750G | 1/1 000 |
| Anti-Phospho-Tau (Ser396) | pS396 | pS396 | H/M | poly | rabbit | Invitrogen #44-752G | 1/10 000 |
| Anti-Phospho-Tau (Ser404) | pS404 | pS404 | M | poly | rabbit | Invitrogen #44-758G | 1/10 000 |
| Anti-total Tau (C-ter ) | Cter | Cter last 15 aa of COOH terminus | M | poly | rabbit | Home-made | 1/10 000 |
| Anti-GAPDH | GAPDH | Mouse GAPDH FL1-335 | H/M | poly | rabbit | Sigma-Aldrich #G9545 | 1/10 000 |

Neuroinflammatory markers were studied using quantitative PCR. Total RNA was extracted from hippocampi and purified using the RNeasyLipid Tissue Mini Kit (Qiagen). One microgram of total RNA was reverse-transcribed using the HighCapacity cDNA reverse transcription kit (Applied Biosystems). Quantitative real-time polymerase chain reaction (qPCR) analysis was performed on an Applied Biosystems™ StepOnePlus™ Real-Time PCR Systems using Power SYBRGreen PCR Master Mix (Applied Biosystems) or TaqMan™ Gene Expression Master Mix (Applied Biosystems™). The thermal cycler conditions were as follows: 95°C for 10 minutes, then 40 cycles at 95°C for 15 seconds and 60°C for 25 seconds for SYBRGreen; and 95°C for 10min, then 40 cycles at 95°C for 15 seconds and 60°C for 1 minute for Taqman. Sequences of primers used in this study are given in Table 2. Cyclophilin A was used as a reference housekeeping gene for normalization. Amplifications were carried out in duplicate and the relative expression of target genes was determined by the ΔΔCt method.

**Table 2.**
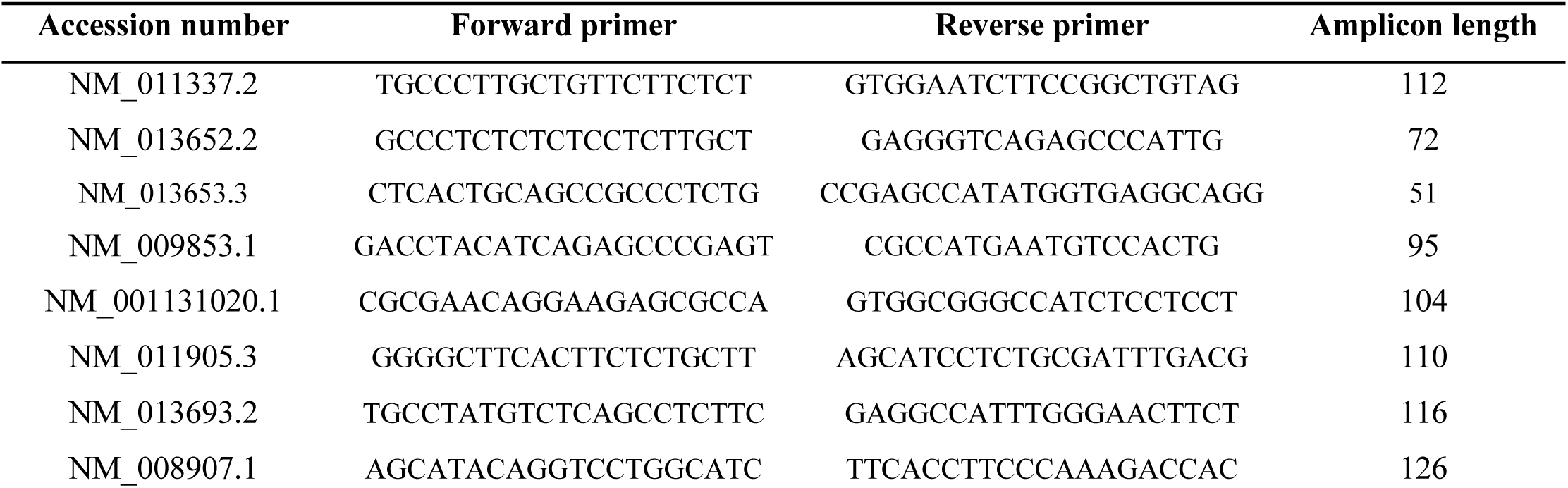

## Results

### Increased synaptic drive in Tau water as compared to Wild type control mice at 8 months

The excitatory and inhibitory synaptic drives received by CA1 pyramidal cells have not been characterized in Tau transgenic mice. We first characterized them in comparison with littermate wild type mice. We found that the frequency of sEPSCs and sIPSCs was increased by 34% (*P* = 0.0285) and 98% (*P* = 0.0135), respectively (Fig. 1 A, E), in Tau water (n= 9 cells, 9 slices, from 5 mice) as compared to wild type controls (n= 8 cells, 8 slices, from 5 mice). The distribution of sEPSC amplitudes was not different but sIPSCs showed a slight (5%) but significant (*P* = 0.0464) increase in amplitude in Tau mice as compared to wild type controls (Fig. 1 B, F). Since spontaneous currents are a mixture of action potential-dependent and independent (miniature) events, we measured miniature currents in the same cells in the presence of TTX (1 µM) to block action potentials. We found that the frequency of miniature EPSCs (mEPSCs) and IPSCs (mIPSCs) was increased by 68% (*P* = 0.0270) and 44% (*P* = 0.0263), respectively (Fig. 2 A, E), in Tau water (n= 9 cells, 9 slices, from 5 mice) as compared to wild type controls (n= 8 cells, 8 slices, from 5 mice). Although the amplitude of mIPSCs was not modified in Tau mice, the amplitude of mEPSCs was increased by 47% (*P* = 0.0253) as compared to Wild type controls (Fig. 2 B, F). The overall synaptic drive received by a neuron is a combination of action potential dependent and independent EPSCs and IPSCs. In order to assess their respective contributions, we calculated the frequency ratio for miniature/spontaneous and EPSC/IPSC events in each recorded cell. In Wild type controls the contribution of miniature events was around 20% and the contribution of EPSCs vs IPSCs was also around 20%, ratios that were not statistically different in Tau water (Fig. 3 A-D). At eight months, the data suggests that there is a general increase in the overall synaptic drive received by CA1 pyramidal cells in Tau mice, whilst both miniature/spontaneous and EPSC/IPSC ratios are not modified.

**Figure 1:**
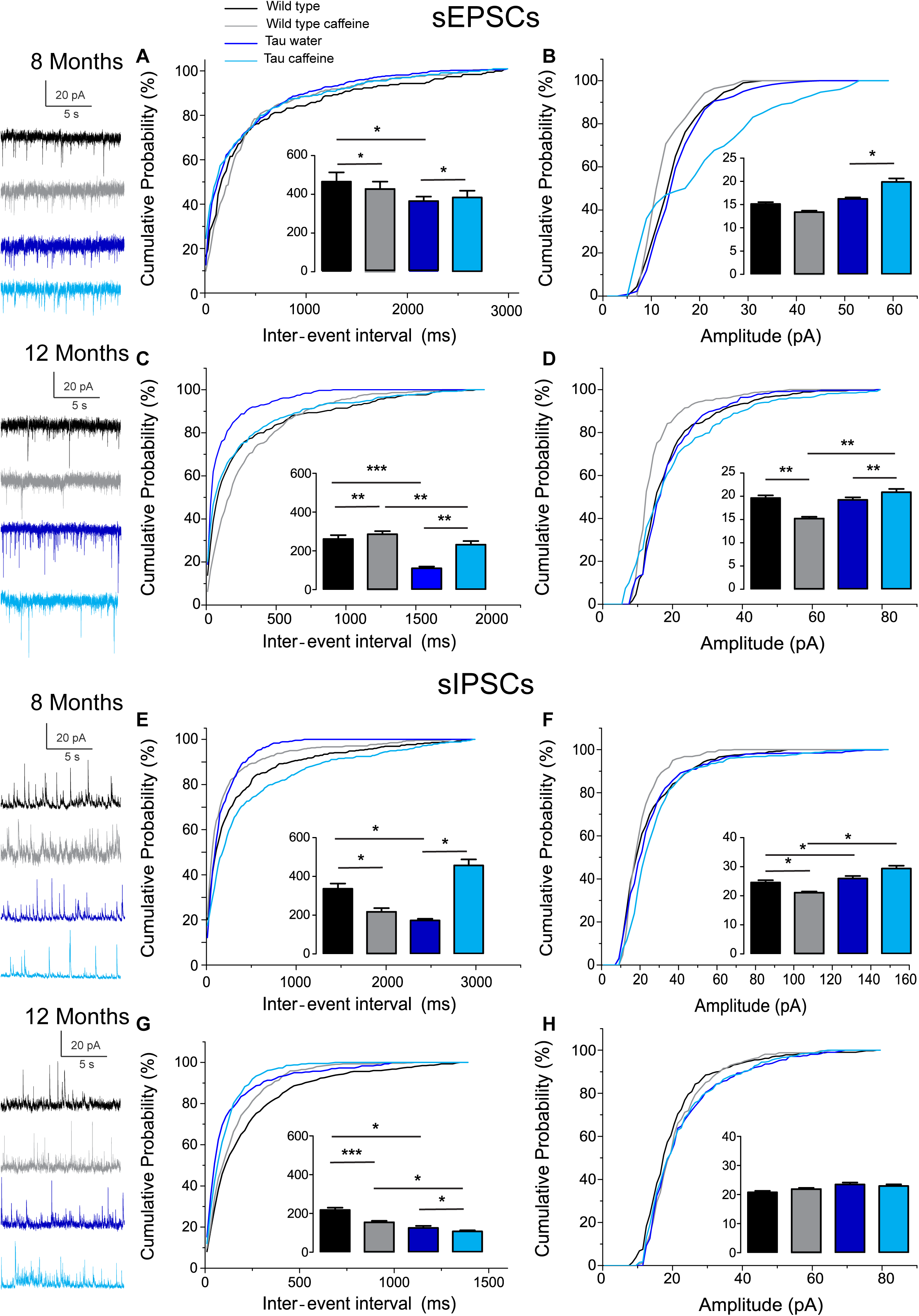
Investigation of chronic caffeine exposure on spontaneous glutamatergic and GABAergic *in vitro* activities in 8 and 12 month-old Wild type and Tau mice. Representative traces of sEPSCs and sIPSCs recorded in CA1 hippocampal pyramidal cells of Wild type water (black), Wild type caffeine-treated (gray), Tau water (blue) and Tau caffeine-treated (light blue) mice are depicted on the left of the cumulative probability histograms (8 month-old sEPSCs: **A, B**; 12 month-old sEPSCs: **C, D;** 8 month-old sIPSCs: **E, F**; 12 month-old sIPSCs: **G, H**). Comparison between cumulative probability distributions was made using the Kolmogorov-Smirnov test (* *P*<0.05; ** *P*<0.01; *** *P*<0.001); bar graphs represent mean ± standard error. n= 8 cells, 8 slices, from 5, 8 month-old Wild type water mice vs. n= 8 cells, 8 slices, from 5, 8 month-old Wild type caffeine-treated mice; n= 8 cells, 8 slices, from 5, 8 month-old Wild type water mice vs. n= 9 cells, 9 slices, from 5, 8 month-old Tau water mice; n= 9 cells, 9 slices, from 5, 8 month-old Tau water mice vs. n= 9 cells, 9 slices, from 5, 8 month-old Tau caffeine-treated mice; n= 8 cells, 8 slices, from 5, 8 month-old Wild type caffeine-treated mice vs. n= 9 cells, 9 slices, from 5, 8 month-old Tau caffeine-treated mice; n= 8 cells, 8 slices, from 5, 12 month-old Wild type water mice vs. n= 8 cells, 8 slices, from 5, 12 month-old Wild type caffeine-treated mice; n= 8 cells, 8 slices, from 5, 12 month-old Wild type water mice vs. n= 7 cells, 7 slices, from 4, 12 month-old Tau water mice; n= 7 cells, 7 slices, from 4, 12 month-old Tau water mice vs. n= 9 cells, 9 slices, from 5, 12 month-old Tau caffeine-treated mice; n= 8 cells, 8 slices, from 5, 12 month-old Wild type caffeine-treated mice vs. n= 9 cells, 9 slices, from 5, 12 month-old Tau caffeine-treated mice.

**Figure 2:**
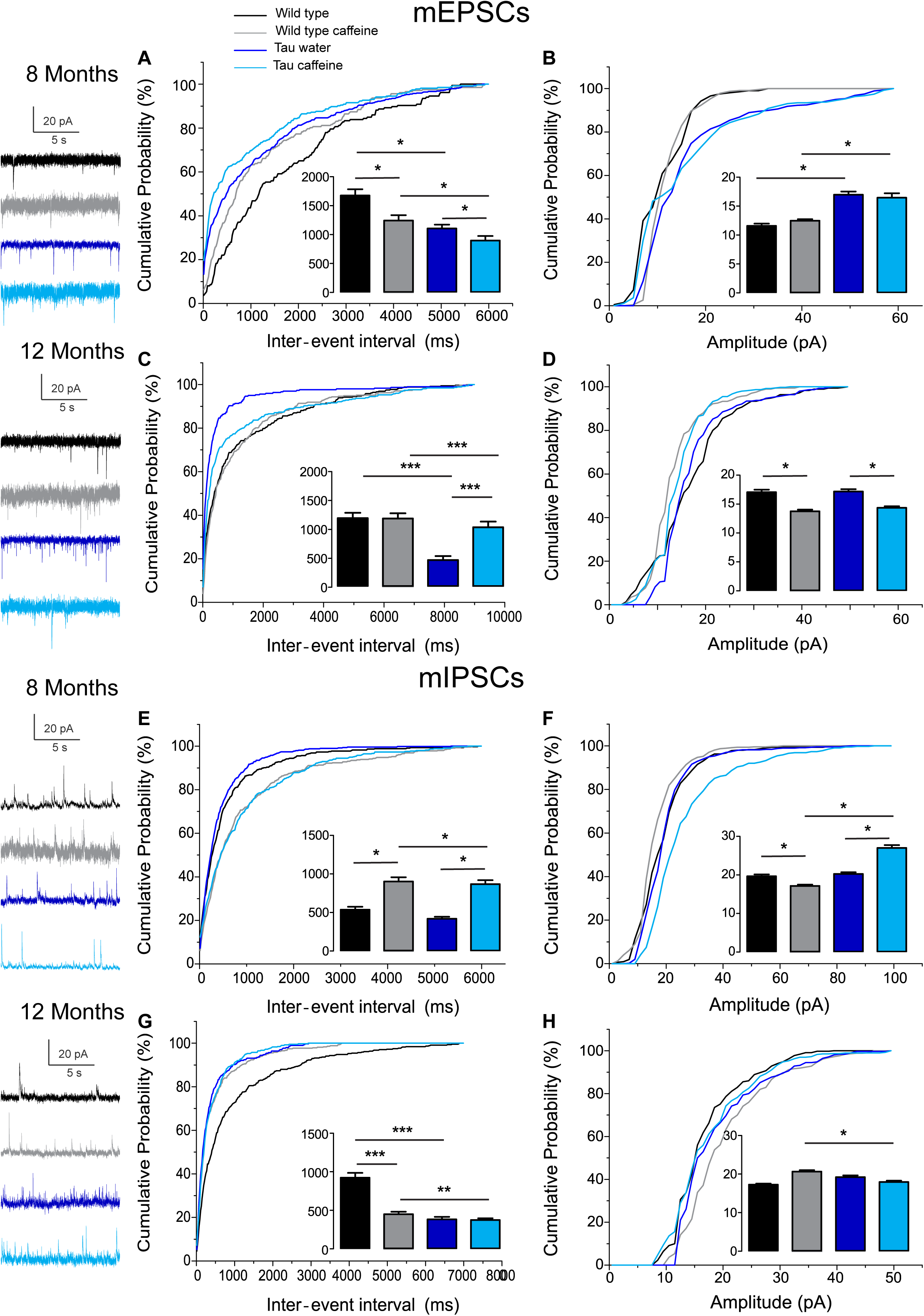
Investigation of chronic caffeine exposure on miniature glutamatergic and GABAergic *in vitro* activities in 8 and 12 month-old Wild type and Tau mice. Representative traces of mEPSCs and mIPSCs recorded in CA1 hippocampal pyramidal cells of Wild type water (black), Wild type caffeine-treated (gray), Tau water (blue) and Tau caffeine-treated (light blue) mice are depicted on the left of the cumulative probability histograms (8 month-old mEPSCs: **A, B**; 12 month-old mEPSCs: **C, D;** 8 month-old mIPSCs: **E, F**; 12 month-old mIPSCs: **G, H**). Comparison between cumulative probability distributions was made using the Kolmogorov-Smirnov test (* *P*<0.05; ** *P*<0.01; *** *P*<0.001); bar graphs represent mean ± standard error. n= 8 cells, 8 slices, from 5, 8 month-old Wild type water mice vs. n= 8 cells, 8 slices, from 5, 8 month-old Wild type caffeine-treated mice; n= 8 cells, 8 slices, from 5, 8 month-old Wild type water mice vs. n= 9 cells, 9 slices, from 5, 8 month-old Tau water mice; n= 9 cells, 9 slices, from 5, 8 month-old Tau water mice vs. n= 9 cells, 9 slices, from 5, 8 month-old Tau caffeine-treated mice; n= 8 cells, 8 slices, from 5, 8 month-old Wild type caffeine-treated mice vs. n= 9 cells, 9 slices, from 5, 8 month-old Tau caffeine-treated mice; n= 8 cells, 8 slices, from 5, 12 month-old Wild type water mice vs. n= 8 cells, 8 slices, from 5, 12 month-old Wild type caffeine-treated mice; n= 8 cells, 8 slices, from 5, 12 month-old Wild type water mice vs. n= 7 cells, 7 slices, from 4, 12 month-old Tau water mice; n= 7 cells, 7 slices, from 4, 12 month-old Tau water mice vs. n= 9 cells, 9 slices, from 5, 12 month-old Tau caffeine-treated mice; n= 8 cells, 8 slices, from 5, 12 month-old Wild type caffeine-treated mice vs. n= 9 cells, 9 slices, from 5, 12 month-old Tau caffeine-treated mice.

**Figure 3:**
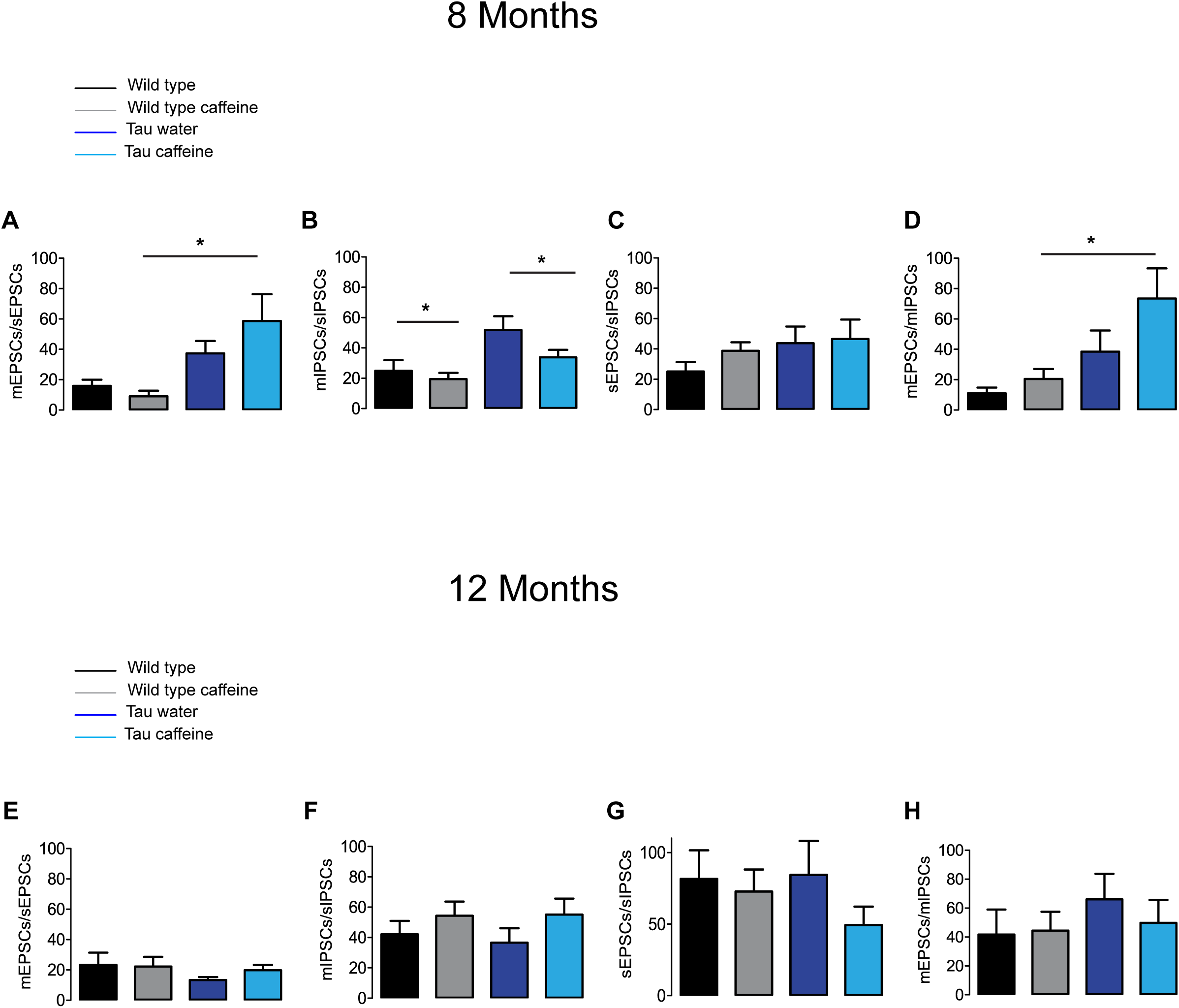
Investigation of chronic caffeine exposure on miniature/spontaneous EPSC/IPSC ratios in 8 and 12 month-old mice. The ratios of the frequencies of mEPSCs to sEPSCs (**A, E**), mIPSCs to sIPSCs (**B, F**), sEPSCs to sIPSCs (**C, G**) and mEPSCs to mIPSCs (**D, H**) were analyzed for Wild type water (black), Wild type caffeine-treated (gray), Tau water (blue) and Tau caffeine-treated (light blue) 8 and 12 month-old mice. Bar graphs represent mean ± standard error; comparison between ratios was made using One way Anova-TukeyKramer post doc test (* *P*<0.05) n= 8 cells, 8 slices, from 5, 8 month-old Wild type water mice vs. n= 8 cells, 8 slices, from 5, 8 month-old Wild type caffeine-treated mice; n= 9 cells, 9 slices, from 5, 8 month-old Tau water mice vs. n= 9 cells, 9 slices, from 5, 8 month-old Tau caffeine-treated mice; n= 8 cells, 8 slices, from 5, 8 month-old Wild type caffeine-treated mice vs. n= 9 cells, 9 slices, from 5, 8 month-old Tau caffeine-treated mice; n= 8 cells, 8 slices, from 5, 12 month-old Wild type water mice vs. n= 8 cells, 8 slices, from 5, 12 month-old Wild type caffeine-treated mice; n= 9 cells, 9 slices, from 5, 12 month-old Tau water mice vs. n= 9 cells, 9 slices, from 5, 12 month-old Tau caffeine-treated mice; n= 8 cells, 8 slices, from 5, 12 month-old Wild type caffeine-treated mice vs. n= 9 cells, 9 slices, from 5, 12 month-old Tau caffeine-treated mice.

### Effects of early life caffeine exposure in Wild type mice evaluated at 8 months

We found that the frequency of sEPSCs and sIPSCs was increased by 17% (*P*= 0.0385) and 54% (*P* = 0.0144, respectively (Fig. 1 A, E), in Wild type caffeine (n= 8 cells, 8 slices, from 5 mice) as compared to wild type controls (n= 8 cells, 8 slices, from 5 mice). Caffeine exposure did not affect the amplitude of sEPSCs but decreased the amplitude of sIPSCs by 14% (*P =* 0.0262) (Fig. 1 B, F). The frequency of mEPSCs was increased by 42% (*P =* 0.0157) whilst the frequency of mIPSCs was decreased by 41% (*P* = 0.0169) as compared to Wild type (Fig. 2 A, E). As for sEPSCs and sIPSCs, caffeine did not affect the amplitude of mEPSCs, but decreased mIPSC amplitude by 12.8% (*P =* 0.0216) as compared to Wild type (Fig. 2 B, F). Caffeine treatment did not affect the miniature/spontaneous and EPSC/IPSC ratios (Fig. 3 A-D). Thus, as reported in 3 months-old offspring of caffeine-exposed mice, there is an hyperactivity of GABAergic networks (Silva CG et al., 2013). However, we did not find hypoactivity of glutamatergic networks found in 3 months-old offspring (Silva CG et al., 2013), perhaps reflecting the time-dependent reorganization of hippocampal networks.

### Effects of early life caffeine exposure in Tau mice evaluated at 8 months

Tau mice exposed to caffeine during gestation and lactation (n= 9 cells, 9 slices, from 5 mice) showed a small decrease in sEPSC frequency by 10% (*P* = 0,0244) and a large decrease in sIPSC frequency by 62% (*P* = 0.0165) as compared to Tau water mice (Fig. 1 A, E). Therefore, although caffeine exposure resulted in increase of sIPSC frequency in wild type mice, it produced a decrease in Tau mice. Although caffeine exposure results in a decreased amplitude of sIPSCs in Wild type mice, the amplitude was not changed in Tau mice (Fig. 1 F). In contrast, the amplitude of sEPSCs was increased by 23% (*P* = 0.0408) in Tau caffeine mice as compared to Tau water mice (Fig. 1 B). As compared to the Tau water group, Tau mice exposed to caffeine during gestation and lactation showed a 38% (*P*= 0.0135) increase in mEPSC frequency and a decrease in mIPSCs frequency by 52% (*P* = 0.0256) (Fig. 2 A, E). In Tau caffeine, the amplitude of mEPSCs was not modified but the amplitude of mIPSCs was increased by 14% (*P* = 0.0285) as compared to Tau water mice (Fig. 2 B, F). Caffeine treatment did not affect the miniature/spontaneous and EPSC/IPSC ratios (Fig. 3 A-D).

### Effects of early life caffeine exposure in Wild type mice evaluated at 12 months

Although early life exposure to caffeine resulted in a slight increase of sEPSC frequency at 8 months, it produced a decrease of sEPSC frequency by 19% (*P* = 0.0025) as compared to Wild type on water at 12 months (Fig. 1 C). The frequency of sIPSCs was increased by 59% (*P* = 1.1151^e-8^) in Wild type caffeine mice, similarly to what we observed at 8 months. Caffeine treatment did not affect the amplitudes of sIPSCs but produced a decrease of sEPSC amplitude by 22% (*P* = 0.0026) (Fig. 1 D, H). Caffeine did not affect mEPSC frequency but increased mIPSC frequency by 106% (*P* = 8.2161^e-6^) as compared to Wild type on water (Fig. 2 C, G), in contrast to that observed at 8 months. Caffeine exposure resulted in a decrease of mEPSC amplitude by 19% (*P* 0,0391) but did not affect that of mIPSCs (Fig. 2 D, H). Therefore the most striking effect of early life exposure to caffeine in Wild type animals is the age-dependent large increase in GABAergic activity received by CA1 pyramidal cells.

### Effects of early life caffeine exposure in Tau mice evaluated at 12 months

In Tau caffeine mice (n= 9 cells, 9 slices, from 5 mice) we found a large decrease in sEPSC frequency by 53% (*P* = 0.0013) as compared to Tau on water mice (Fig. 1 C), in keeping with the effect of caffeine on sEPSC frequency described above in wild type mice exposed to caffeine. Although there was an increase in sEPSC at 8 and 12 months in Tau water mice, caffeine treatment prevented such increase. Although caffeine treatment results in a decreased sIPSC frequency at 8 months, we found an increase in sIPSC frequency by 17% (*P* = 0.0216) at 12 months as compared to Tau on water (Fig. 1 G). Caffeine exposure resulted in a slight increase of sEPSC amplitude by 9% (*P*= 0.0191) but did not change sIPSC amplitude as compared to Tau on water (Fig. 1 C, G). In Tau mice exposed to caffeine there was a large decrease in mEPSC frequency by 55% (*P* = 4.99691^e-50^), while mIPSC frequency was not modified (Fig. 2 D, H). The amplitude of mEPSCs (but not that of mIPSCs) was decreased by 16% (*P* = 0.0391) as compared to Tau on water (Fig. 2 D, H). Thus, during aging, early exposure to caffeine exacerbates the increase in GABAergic drive received by CA1 pyramidal cells in Tau mice.

### Reorganization during aging: differences between 8 month- and 12 month-old Wild type and Tau mice on water

Since our hypothesis is that early-life exposure to caffeine is accelerating the occurrence of phenotypic traits, we now compare the most striking modifications between 8 month- and 12 month-old animals. In both Wild type (n= 8 cells, 8 slices, from 5 mice) and Tau mice on water (n= 7 cells, 7 slices, from 4 mice) we found a large increase in sEPSC frequency by 87% and 231%, respectively, as well as in sIPSC frequency by 55% and 38%, respectively (Fig. 1 A, C, E, G). The amplitude of sEPSCs increased by 30% and 19 %, respectively, in Wild type and Tau mice on water, while for sIPSCs we found a decrease by 20% and 26%, respectively (Fig. 1 B, D, F, H). The frequency of mEPSCs also increased at 8 months in both strains (81% and 72% respectively), but the frequency of mIPSCs decreased (42%) in Wild type whilst it increased (10%) in Tau mice (Fig. 2 A, C, E, G). The amplitude of mEPSCs increased by 47% and 1% in Wild type and Tau mice on water, respectively, while for mIPSCs it decreased by 13% and 5%, respectively (Fig. 2 B, D, F, H). During aging, at least for the two time points considered in this study, there is a global increase of glutamatergic and GABAergic activity received in CA1 pyramidal cells. Interestingly, the fraction of action potential-dependent events is largely increased in 12 month-old wild type as compared to 8 months (since the contribution of mIPSCs is decreased).

### Effects of early-life exposure to caffeine on learning and memory performance

Hippocampal dependent memory was assessed using the Barnes maze test (Koopmans et al., 2003) in 8 month-old mice (n= 11 wild type water, 13 wild type caffeine-treated, 11 TAU water, 13 TAU caffeine-treated mice). Training was performed for 3 days (D1, D2, D3), with 3 trials (T1, T2, T3) per day. We found the typical learning curve in wild type mice on water as assessed by the latency to the target (Fig. 4 G). The learning curve was similar in Wild type caffeine mice, while a delay in learning was found in both Tau groups (treated or not with caffeine) with a significant difference at D1T2 and D2T1 between Wild Type and Tau mice (Fig. 4 G). No difference between was found from D2T2. Thus, caffeine exposure by itself does not affect the learning curve, but the underlying Tau pathology does.

One week after the learning part, we performed a probe test (P1) and a second one, one week later (P2) to assess memory retention. The caffeine exposed Tau group showed a significant decrease in the number of holes searched in the GOAL area during P2 as compared to P1 (56 %, *P* = 0.0383), which shows an impairment to remember the general location of the escape hole in Tau mice exposed to caffeine (Fig. 4 A). The latency to the escape hole was increased by 181% (*P =* 0.0003) in Tau caffeine mice at P2 (Fig. 4 H). The same trends were found in Wild type animals exposed to caffeine but it was not significant. Thus, caffeine exposed Tau mice show memory deficits.

**Figure 4:**
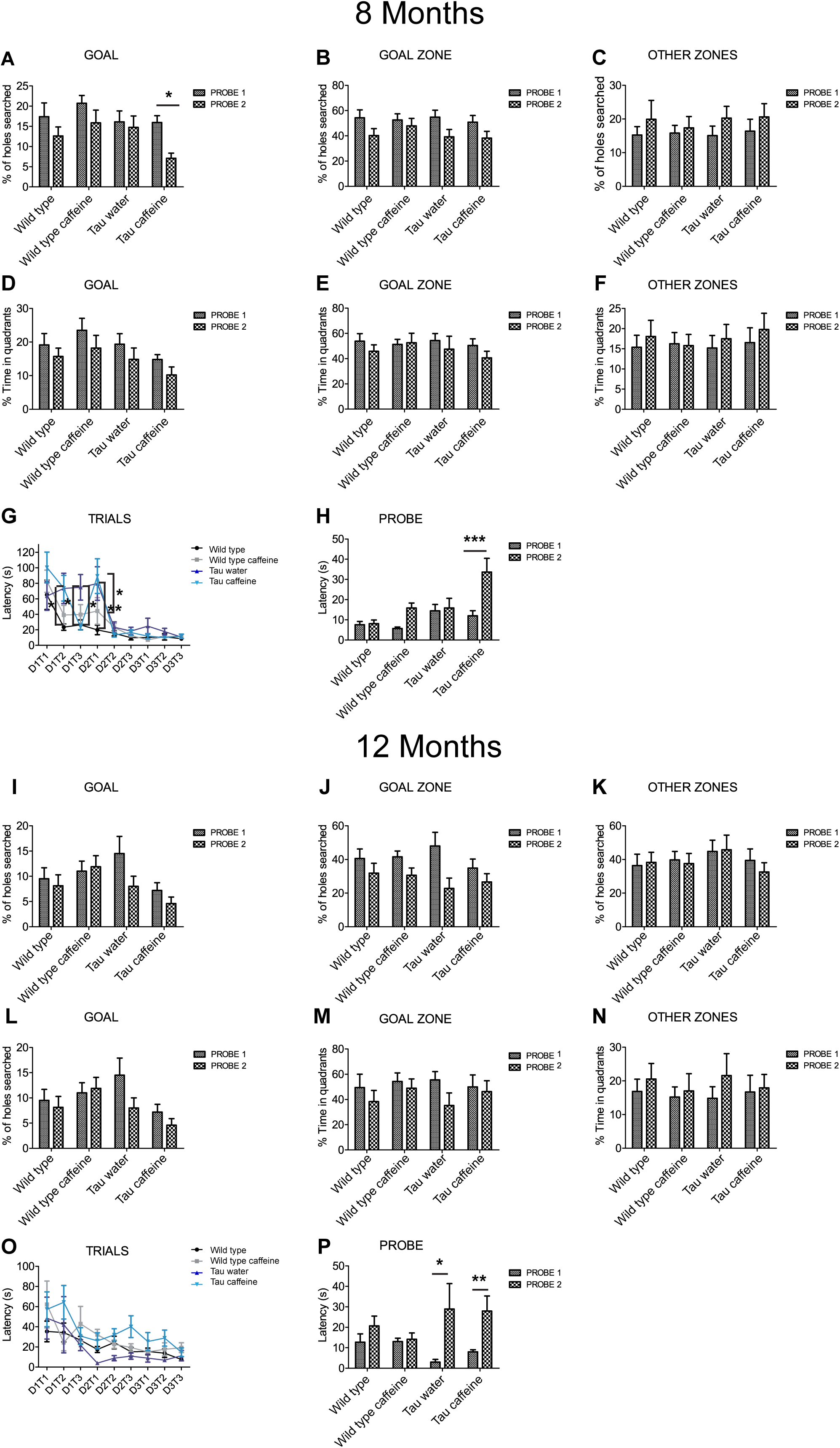
Investigation of early-life exposure to caffeine on learning and memory performance. Barnes maze test was performed in 8 **(A-G)** and 12 **(I-O)** month-old mice. Training was performed for 3 days (D1, D2, D3), with 3 trials (T1, T2, T3) per day. For 8 and 12 month-old mice, the % of holes searched in GOAL **(A, I)**, GOAL ZONE **(B, J)** and OTHER ZONES **(C, K)**, the % time in quadrants **(D, L)**, GOAL ZONE **(E, M)** and OTHER ZONES **(F, N)** and the latency **(G, O)** were analyzed. To assess memory retention one week after the learning part, a probe test (P1) was performed and a second one, one week later (P2) on 8 and 12 month-old mice **(H, P)**. Two way Anova followed by Bonferroni post hoc test (* *P*<0.05; ** *P*<0.01; *** *P*<0.001). n= 11 Wild type water 8 month-old mice vs 11 Tau water 8 month-old mice at D1T2; n= 11 Wild type water 8 month-old mice vs 11 Tau water 8 month-old mice at D2T1; n= 11 Tau water 8 month-old mice vs 11 Tau caffeine-treated 8 month-old mice at D1T3; n= 11 Tau water 8 month-old mice vs 11 Tau caffeine-treated 8 month-old mice at D2T1; n = 13 Tau caffeine-treated 8 month-old mice at P1 vs n = 13 Tau caffeine-treated 8 month-old mice at P2; n = 4 Tau water 12 month-old mice at P1 vs n = 4 Tau water 12 month-old mice at P2; n = 10 Tau caffeine-treated 12 month-old mice at P1 vs n = 10 Tau caffeine-treated 12 month-old mice at P2.

We performed the same study in another series of 12 month-old animals (n= 8 wild type water, 10 wild type caffeine-treated, 4 TAU water, 10 TAU caffeine-treated mice). Surprisingly, in contrast to 8 month-old animals, the learning curves were similar in the four groups, although the TAU caffeine group showed a tendency for slower learning. Both Tau water and Tau caffeine groups showed a decrease in the number of holes searched in the GOAL area at P2, by 45% and 36% respectively, but the decrease was not significant. However, the latency at P2 showed an increase by 877.5% (*P =* 0.0122) and 249.9% (*P =* 0.0012) for Tau water and Tau caffeine, respectively (Fig. 4 P), confirming an impairment in memory retention for Tau mice exposed to an early caffeine consumption and the appearance of the same deficit in the Tau control group. This suggests that caffeine exposure in Tau mice enabled an earlier expression of memory deficits at 8 months.

### Impact of early life caffeine consumption on hippocampal Tau phosphorylation and related neuroinflammatory markers

THY-Tau22 mice exhibit progressive memory impairments in parallel with the development of hippocampal Tau hyperphosphorylation and neuroinflammation (Van der Jeugd et al., 2013; Laurent et al., 2017). At 6-8 months of age, Tau pathology and neuroinflammation are ongoing in the hippocampus of THY-Tau22 mice. The potentiation of memory deficits by early-life caffeine in Tau mice opened the possibility that Tau pathology itself also associated neuroinflammation might have been advanced, therefore we performed biochemical and qPCR experiments in an additional group of animals at the age of 6 months. Using sodium dodecyl sulphate-polyacrylamide gel electrophoresis, we evaluated Tau phosphorylation in both Tau experimental groups using antibodies raised against several Tau phosphoepitopes. None of the epitope studied was modified by early life exposure to caffeine (Fig. 5A). As caffeine modulates neuroinflammation (Brothers et al., 2010; Laurent et al., 2014), we checked several neuroinflammatory markers previously described to be early or lately upregulated in the hippocampus of THY-Tau22 animals (Laurent et al., 2017). In general, in accordance with the absence of impact on Tau phosphorylation, early caffeine treatment did not modulate any of the neuroinflammatory markers studied. We could only evidence a slight reduction of the expression of the microglia CD68 markers in TAU mice treated with caffeine vs. Tau water animals (Fig. 5B).

**Figure 5:**
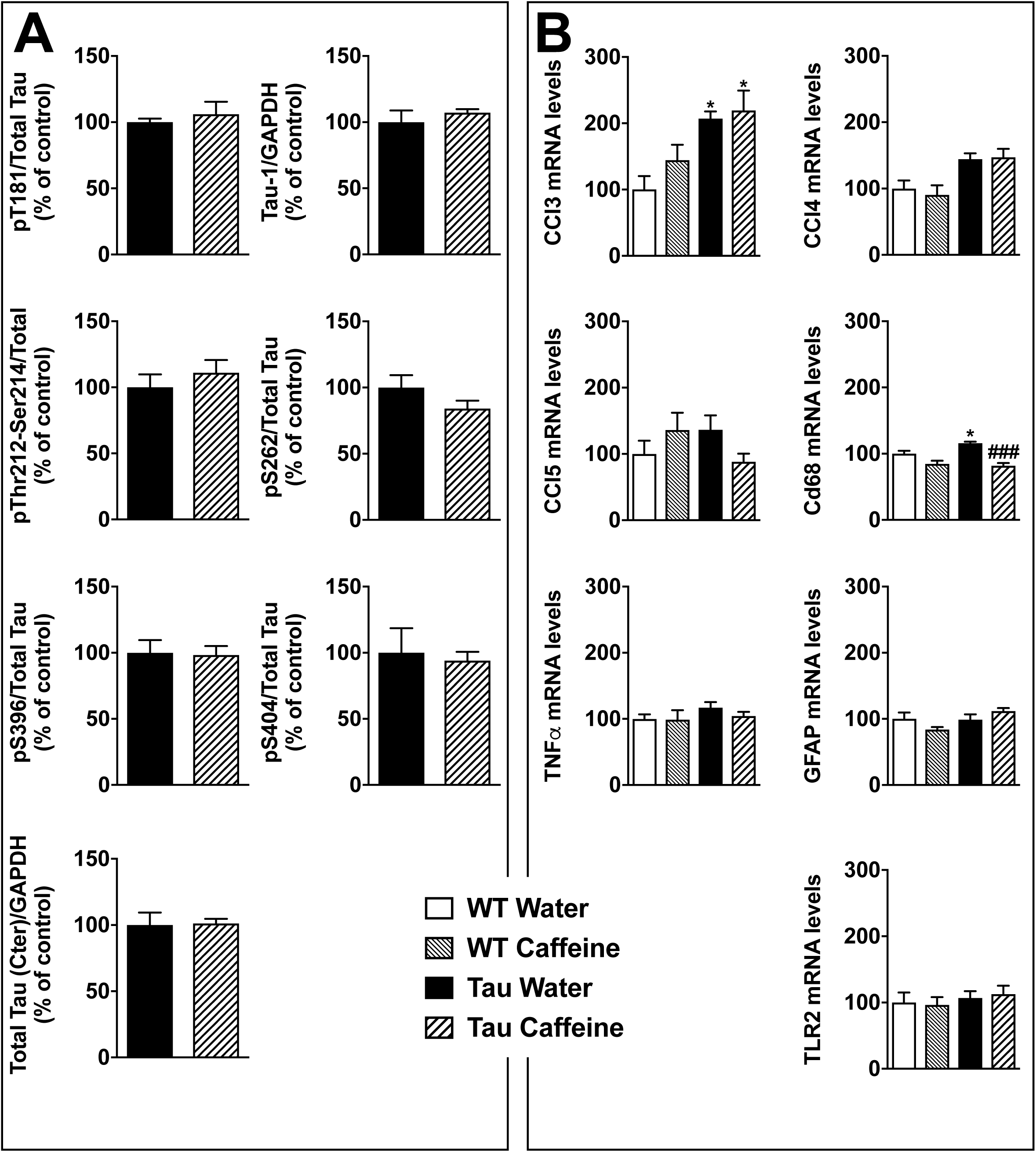
Investigation of early-life exposure to caffeine on hippocampal Tau phosphorylation and associated neuroinflammatory markers. (A) Western blot analysis of tau phosphorylation in water or caffeine exposed THY-Tau22 offsprings using antibodies targeting physiological (pThr181, Tau-1, pSer262, pSer396, pSer404) and pathologic (pThr212/Ser214) Tau epitopes. Quantifications were performed over total tau levels (Cter). Total tau levels were quantified versus GAPDH, used as loading control (n=7/group). (B) qPCR analysis of hippocampal neuroinflammatory markers associated with Tau pathology in the THY-Tau22 model (n=6-10 per group).

## Discussion

The most salient feature of the results is the occurrence of cognitive deficits already at 8 months in the Barnes maze in caffeine exposed Tau mice. These deficits occurred at 12 months in water exposed Tau mice and were still present at that age in caffeine Tau mice. In contrast, caffeine treatment did not affect cognitive performance in Wild type mice. Using different tests, we previously reported cognitive deficits in caffeine-exposed Wild type mice at 6 months (Silva et al., 2013). The discrepancy may stem from the use of different tests and different strains: C57BL6/J here, and GIN FVB/N mice in (Silva et al., 2013). In the present work, we focused on the Barnes maze, which is a terrestrial version of the Morris water maze. We did not use other tests to prevent interference when using different behavioral tests in succession. Humans have different genetic backgrounds and go through different life experiences, which will determine their sensitivity to the development of diseases, in particular neurological disorders. The fact that caffeine exposure during early life produces an earlier occurrence of cognitive deficits in the Tau model used here may be relevant to a subset of human individuals. Said differently, exposure to caffeine during pregnancy may sensitize some but not all offsprings. Future studies will need to investigate other time points, other tests and other models of AD. The experimental protocol is however highly time-consuming. We cannot rely on established aging colonies, since females need to be exposed to caffeine in the drinking water before mating until weaning. Then, it is necessary to wait until offspring reach the appropriate age.

Although caffeine exposure did result in earlier cognitive deficits in Tau mice, it is difficult to propose a mechanism. Previous works in control GIN (GFP-expressing Inhibitory Neurons) mice have shown that caffeine slows down the migration of GABA neurons resulting in a delayed insertion in the circuitry, both in the hippocampus and the cortex during the first postnatal week (Silva et al., 2013; Fazeli et al., 2017). This mechanism of age-dependent refinement of local circuit inhibition may have a direct impact on the hippocampal network activity and performance during development (Salesse, 2011). Such morphological changes are associated with enhanced sensitivity to epilepsy and hyperactivity *in vivo* (Silva et al., 2013; Fazeli et al., 2017). Hence, caffeine exposure has important physiological consequences, which could be widespread in the body. The cognitive deficits constitute a read-out, but multiple reasons could explain them (activation of the HPA axis, inflammation, metabolic defect, cell death, synaptopathy, channelopathy, epigenetic modifications, to name but a few). In THY-Tau22 mice, cognitive deficits have been in general associated with Tau pathological load and associated neuroinflammation (Van der Jeugd et al., 2013; Laurent et al., 2016; Laurent et al., 2017, but see Chatterjee et al., 2018 and Burlot et al., 2015). However, analysis of cardinal pTau and neuroinflammatory markers did not correlate with impact of gestational caffeine on memory abilities of offsprings. As another physiological readout, we focused on the GABAergic and glutamatergic drives received by CA1 pyramidal cells *in vitro*. This provides a general view of how the CA1 network may be reorganized. It is important to note that, although the Barnes maze test involves the hippocampus, the information coming from the analysis in CA1 cannot be used to explain cognitive deficits in a causal manner, it can only be be correlative. In addition, data interpretation is difficult because of the presence of three independent variables imposed by the experimental protocol. The first two independent variables are caffeine and TAU, both of which result in morpho-physiological alterations. The third independent variable is age, since we are considering two time points, 8 and 12 months. Yet, the *in vitro* approach provides interesting results in their complexity. Since our hypothesis is that caffeine exposure accelerates the occurrence of the phenotype, we start to discuss the age factor. In WT mice on water, we found an increase in both glutamatergic and GABAergic drives between 8 and 12 months. We can speculate that during normal aging, there is a gradual increase of both excitatory and inhibitory inputs received by CA1 pyramidal cells. More time points should be investigated to test this hypothesis. This result is consistent with that reported in the prefrontal cortex during aging (using later time points) for animals preserving their cognitive performances (Bories et al., 2013). Interestingly, in Tau mice on water, both glutamatergic and GABAergic drives were increased at 8 and 12 months as compared to their Wild type counterparts. In keeping with our results, most studies using Tau or Aß models report an increase in the excitatory drive as compared to Wild Type (Crimins et al., 2011; Dalby et al., 2014; Ovsepian et al., 2017), but see (Rocher et al., 2008). Less data is available for the GABAergic drive. In the cortex of a different Tau model, there was no difference in sIPSC frequency at 9 months as compared to Wild Type (Crimins et al., 2011). The discrepancy with our results may stem from the type of mutant used and the brain region selected. Together, our results raise the intriguing possibility that the Tau mutation accelerates the aging process as assessed with the glutamatergic and GABAergic drives received by CA1 pyramidal cells.

Early life exposure to caffeine changed the apparent co-variance relationship between the two age and Tau independent variables. At 8 months, both GABAergic and glutamatergic drives are decreased in Tau caffeine mice as compared to Tau water mice. Whether this decrease is related to the early occurrence of deficits in the Barnes maze in not known. However, at 12 months, although the excitatory drive is decreased as compared to that measured in Tau water mice, the inhibitory drive is increased. This exemplifies the difficulty in interpreting the *in vitro* data. However, our data emphasize that caffeine exposure disrupts the effect of the pathogenic process characteristic of the Tau phenotype in CA1 pyramidal cells.

In conclusion, we propose that early caffeine exposure produces physiological and cognitive alterations in a Tau pathological context, supporting our hypothesis that caffeine consumption during pregnancy may constitute a risk-factor for an earlier development of AD-related phenotypes. Future studies are needed using other mouse models, and, as importantly, to determine whether re-exposure to caffeine during adulthood is protective.

## Acknowledgements

This work was supported by a grant from Association France Alzheimer, project AAP SM 2016/1567. We thank the Animal Facility (F-59000 Lille, France) and Mélanie Besegher, Cyrille Degraeve, Caroline Declerck, Kim Letten, Yann Lepage, Benjamin Guerrin, Didier Montignies, Christian Meunier, Quentin Dekeyser and Romain Dehaynin for animal care. Inserm UMR-S1172 is supported by grants from Hauts-de-France (PARTEN-AIRR, COGNADORA), ANR (ADORATAU and ADOSTAsTRAU to DB; GRAND and TONIC to LB) and Programs d’Investissements d’Avenir LabEx (excellence laboratory) DISTALZ (Development of Innovative Strategies for a Transdisciplinary approach to ALZheimer’s disease), Fondation pour la Recherche Médicale, France Alzheimer/Fondation de France, FHU VasCog research network (Lille, France), Fondation Vaincre Alzheimer, Fondation Plan Alzheimer, Inserm, CNRS, Université Lille, Lille Métropole Communauté Urbaine, DN2M.

